# A new dispersal-informed null model for community ecology shows strong performance

**DOI:** 10.1101/046524

**Authors:** Eliot Miller

## Abstract

Null models in ecology have been developed that, by maintaining some aspects of observed communities and repeatedly randomizing others, allow researchers to test for the action of community assembly processes like habitat filtering and competitive exclusion. Such processes are often detected using phylogenetic community structure metrics. When biologically significant elements, such as the number of species per assemblage, break down during randomizations, it can lead to high error rates. Realistic dispersal probabilities are often neglected during randomization, and existing models make the oftentimes empirically unreasonable assumption that all species are equally probable of dispersing to a given site. When this assumption is unwarranted, null models need to incorporate dispersal probabilities. I do so here, and present a dispersal null model (DNM) that strictly maintains species richness, and approximately maintains species occurrence frequencies and total abundance. I tested its statistical performance when used with a wide breadth of phylogenetic community structure metrics across 3,000 simulated communities assembled according to neutral, habitat filtering, and competitive exclusion processes. The DNM performed well, exhibiting low error rates (both type I and II). I also implemented it in a re-analysis of a large empirical dataset, an abundance matrix of 696 sites and 75 species of Australian Meliphagidae. Although the overall signal from that study remained unchanged, it showed that statistically significant phylogenetic clustering could have been an artifact of dispersal limitations.

## INTRODUCTION

Null models in ecology are often used to test whether an assemblage of co-occurring species differs from what would be expected of a random assortment of co-occurring species (Gotelli and McGill 2006). Of particular interest here has been determining whether a community shows evidence that competition structures its constituent member species. How to define random in the null model context has been a matter of great contention since the debates of the 1970s and 1980s (Connor and Simberloff 1979; Diamond and Gilpin 1982; Connor and Simberloff 1983). Ideally, null models shuffle elements of the observed data related to the null hypothesis and preserve unrelated aspects of the observed data (e.g., which vs. how many species co-occur). Null models often take the form of repeated randomizations of an observed community data matrix (CDM), which are then compared to the observed CDM to detect non-random patterns of community assembly. Matters of contention include what elements of the CDM should be maintained (e.g., row and/or column sums, Diamond and Gilpin 1982), the most efficient algorithms for shuffling matrices (Miklós and Podani 2004), what metrics should be used to calculate patterns in the matrices (Stone and Roberts 1990), and which null models provide the best statistical performance (Gotelli 2000). How to incorporate abundance instead of simply presence-absence into these null models is another important research focus that has to date only received some attention (Hardy 2008; Ulrich and Gotelli 2010; Miller et al. 2016).

Furthering complicating the field has been the rise of neutral models (Bell 2000; Hubbell 2001). Here, I follow Gotelli and McGill (2006), in considering neutral models as best used for testing whether per capita demographic rates differ between species. As the focus of this paper is on testing whether species interactions are important, and assumes that species differ in their per capita demographic rates, this paper is about a new null model. That said, neutral models derive from the theory of island biogeography, which is governed by the countervailing forces of dispersal and extinction. Because this paper introduces a null model that incorporates dispersal, it falls closer to what Gotelli and McGill consider a process-based model that “crosses the line”. This is similar to a recent method incorporating speciation into null expectations (Pigot and Etienne 2015).

In this paper I define a CDM as a matrix with sites (e.g., quadrats, samples, plots) as rows and species as columns. The CDM as defined here can and, because of the additional detail afforded, ideally does incorporate relative or absolute abundance. During the null model debates, the focus was on demonstrating matrix-wide departures from expectations. In other words, the question was whether the entire CDM showed evidence of non-randomness. Accordingly, the metrics used to document the significance, or lack thereof, in community structural patterns were calculated per matrix, and generally dealt with presence-absence matrices (Schluter 1984; Stone and Roberts 1990). With the more recent focus on phylogenetic community structure (Webb 2000), the focus has shifted away from CDM-level patterns to assemblage-level (i.e. row-level) structural patterns. With this has come the introduction of a host of new phylogenetic community structure metrics, many of which incorporate abundance (Faith 1992; Webb 2000; Helmus et al. 2007; Cadotte et al. 2010; Kembel et al. 2010; Miller et al. 2013). These metrics quantify the relatedness of co-occurring species, with the assumption being that closely related co-occurring species provide evidence of habitat filtering, while distantly related co-occurring species provide evidence of competitive exclusion (Webb 2000). The null model introduced in this paper is intended for use in assessing assemblage-level patterns, and its statistical performance was tested here in that context. Its relevance to matrix-level patterns of co-occurrence is not tested here.

Many null models shuffle species’ presences or abundances within rows, allowing species to occur with equal probability in the randomized matrices. Various improvements have been developed, including models that maintain species’ occurrence frequencies (Gotelli 2000), both species richness and occurrence frequency (Miklós and Podani 2004), and elements of species’ abundance distributions (Hardy 2008). Such models have been shown to reduce type I errors (Gotelli 2000; Miller et al. 2016). Miller et al. (2016) showed that a further reduction in error rates can be achieved by creating a CDM *de novo* that mimics regional dispersal pressures on a “local” community (the CDM). However, in all of these null models quadrats are disassociated from their geographic realities. Current null models randomize sites but do not take into account dispersal probabilities between sites in the randomization process. Thus, a species from a distant site is just as likely to be placed in a simulated site as a species from a nearby site. The 3t null model introduced by Hardy (2008) made strides towards addressing this issue, but it does not maintain quadrat species richness and requires transect-like sampling.

Dispersal, even over short distances like those across a forest plot, can greatly influence which species occur where (MacArthur and Wilson 1967; Laurance et al. 2002). Over large distances, such as those between grid cells in the desert of inland Australia and the rainforests of the coast, it can be difficult to parse the influence of community assembly processes such as habitat filtering versus that of physical dispersal limitation. For example, if a certain clade within the study system has diversified within a small region of the continent, significant phylogenetic clustering in that region is not necessarily attributable to habitat filtering, and might be due entirely to a failure of these species to disperse to other regions. A more flexible, dispersal-informed null model is a clear research priority that should prove useful to empirical researchers. Having a null model that respects quadrat-specific dispersal probabilities, while also maintaining quadrat-specific species richness and species’ matrix-wide occurrence probabilities would assist with teasing apart such community assembly subtleties. In this paper, I develop such a null model and test its statistical behavior and performance.

## METHODS

### Description of the dispersal null model

The dispersal null model (DNM) takes as input the original CDM, **C***_O_*, and a symmetrical matrix, **D**, that describes the distances among quadrats. It provides as output a randomized CDM, **C***_R_*, with the same dimensions as **C***_O_*. Distances in **D** can be geographic, climatic, or otherwise of the researcher’s choice. I then define the randomization procedure as follows. Let *i_O_* be a row (quadrat) from **C***_O_*, and *i_R_* be the corresponding row from **C***_R_*. Let *j* be any other row from **C***_O_* where *i_O_* ≠ *j*. Let *SR*(*n*) be the species richness of some row *n* from a CDM. Then, for each *i_O_*, some *j* is sampled with a probability determined by the reciprocal of its distance from *i_O_*. A species is then sampled from *j* with a probability proportional to its abundance in *j*, placed into *i_R_*, and assigned the same abundance as in *j*. Then this process is repeated until *SR*(*i_O_*) = *SR*(*i_R_*). If for any *j*, the species sampled has already been settled into *i_R_*, then it is discarded and another *j* and corresponding species sampled. The model goes on to repeat the process for all quadrats *i*, which results in a filled **C***_R_*. The need to propose and then potentially reject species necessitates the use of a serial loop, which causes the DNM to run more slowly than simpler randomization procedures. I refer to this form of model as DNM_1_.

A slight variation of the model, DNM_2_, generates a normal distribution (with a standard deviation of 1, rounded to whole numbers and values < 1 rounded up to 1) centered on the abundance of the sampled species in *j*. Rather than directly assigning a species the same abundance it had in *j*, a value is sampled from this distribution. This causes a slight slowdown (~5% longer) in null model performance, but theoretically results in additional exploration of null biological space.

A considerable variant of the model (DNM_3_) does not incorporate species’ abundances in *j* into the probability that they will be sampled. Instead, all species present in *j* have an equal probability of being sampled and placed into *i_R_*. Thus a species’ proximity to but not its abundance in *j* influences its probability of settling in *i_R_*. The biological meaning here is changed from DNM_1_. With DNM_3_, any factors influencing individual species’ abundances within a quadrat, such as competition, are less influential in the randomized matrices. While DNM_1_ can help researchers test for non-random patterns of community assembly given realistic dispersal pressures, where both abundance and distance to a focal quadrat matters, DNM_3_ would be more pertinent if the focus was on the influence of competition given distance dispersal limitations only. That is, if a researcher thought it possible that species might be rare in observed quadrats as a consequence of competition, then DNM_3_ would randomize those structures in the observed data and allow that hypothesis to be tested. With DNM_3_, I recommend that researchers assign species abundances by sampling from the vector of observed non-zero abundances in the original CDM.

The three forms of DNM are available in the R package *metricTester* (Miller et al. 2016). Though the DNM runs more slowly than traditional matrix randomization, *metricTester* utilizes multicore processing and thus can manage a reasonable number of randomizations of an observed CDM.

### Statistical behavior and performance of the dispersal null model

As explained above, the DNMs strictly maintain species richness. I was interested in how well they maintain species’ occurrence frequencies and total abundances in the randomized matrices. To test this, I created a CDM with the simulateComm function in *metricTester*. The CDM contained 100 quadrats and species. Species richness varied from 10 to 34, with each value represented four times. Species were assigned abundances by drawing from a log-normal distribution with mean=2 and SD=1 (on a log scale). I then randomized the CDM 20 times with DNM_1_, calculating species’ occurrence frequencies and total abundances after each randomization. I took the mean of these observations and compared it to observed values from the observed CDM. I performed the same procedure but randomized the CDM 20 times with DNM_3_, in this case setting `abundance.assigned` to “overall”.

I used identical methodology as Miller et al. (2016) to test the performance of the DNMs. Appendix S3 of that paper provides schematic illustration of the methodology. Thus, I used the `multiLinker` function with the following parameters. I set `no.taxa` to 100, `arena.length` to √(10^5^), `mean.log.individuals` to 3.5, `length.parameter` to 1000, `sd.parameter` to 40, `max.distance` to 20, `proportion.killed` to 0.2, `competition.iterations` to 60, `no.quadrats` to 20, `quadrat.length` to √(1000), `concat.by` to “both”, and randomizations to 1000. The simulation and performance testing process is as follows. (1) Generate a phylogeny describing the relationships among the species that will be involved in the simulation. (2) Generate realistic spatial arenas of 2,000-4,000 individuals according to either random, habitat filtering or competitive exclusion community assembly rules. In the habitat filtering simulations, individuals are placed in the arena according to spatial preferences that exhibit Brownian motion evolution along the simulated phylogeny. The competitive exclusion simulations begin with the random simulation. Individuals in genetically clustered areas of the arena then compete, resulting in the removal of some of these most closely related individuals. Removed individuals are then replaced according to their initial arena-wide abundances (i.e. simulating regional dispersal pressures), and the entire process is repeated for 60 generations. (3) Create CDMs (one per spatial simulation) by placing 20 quadrats in each arena and determining which individuals fall within each quadrat. (4) Calculate a wide breadth of phylogenetic community structure metrics on the observed quadrats. These metrics were PAE (phylogenetic abundance evenness), *H*_AED_ (community abundance-weighted evolutionary distinctiveness), IAC (imbalance of abundance), *E*_AED_ (equitability abundance-weighted evolutionary distinctiveness), *H*_ED_ (community evolutionary distinctiveness), *E*_ED_ (equitability evolutionary distinctiveness), MNTD (mean nearest taxon distance), AW MNTD (abundance-weighted MNTD), PD (phylogenetic diversity), PD*c* (PD not including the root), MPD (mean pairwise phylogenetic distance), interspecific MPD (interspecific abundance-weighted MPD), intraspecific MPD (intraspecific abundance-weighted MPD), complete MPD (complete abundance-weighted MPD), and PSE (phylogenetic species evenness) (Faith 1992; Webb 2000; Helmus et al. 2007; Cadotte et al. 2010; Kembel et al. 2010; Miller et al. 2013). (5) Use the Euclidean distances between quadrat centroids to randomize each CDM 1,000 times with DNM_1_, and 1,000 times with DNM_3_. Note that this happens across all spatial simulations, such that after the three spatial simulations, then 1,000 randomizations of each observed CDM with each of DNM_1_ and DNM_3_, the result is a collection of 6,000 randomly assembled CDMs and three observed CDMs. (6) After each randomization, calculate all phylogenetic community structure metrics across the assembled CDM and retain. (7) Concatenate the randomized metrics by the quadrat from which they come, and derive a standardized effect score (SES) per observed quadrat as the difference between the observed and the mean of the randomized scores (at that quadrat) divided by the SD of the randomized scores (at that quadrat). (8) Per CDM, use a Wilcoxon signed-rank test to compare the distribution of SES scores to zero. Record a type I error if the distribution differs significantly from zero for the random community assembly, and a type II error if the distribution was not significantly less or greater than zero in the habitat filtering or competitive exclusion simulation, respectively. (9) Repeat the entire process 1,000 times (i.e. run 1,000 each of the neutral, habitat filtering, and competitive exclusion spatial simulations, with the resulting CDMs randomized 1,000 times each). In this way, per metric per null model, I calculated an average type I error rate as the proportion of 1,000 random communities whose SES distribution differed significantly from zero, and an average type II error rate as the mean of the proportion of communities from either the 1,000 habitat filtering or the 1,000 competitive exclusion simulations whose SES distributions did not differ as expected from zero.

### Testing the dispersal null model on an empirical dataset

I was also interested in whether DNM_1_ could be readily applied to an empirical dataset, and what influence it would have on the interpretation of results. Miller et al. (2013) found strong evidence that phylogenetic niche conservatism shapes which lineages of Australian honeyeaters (Meliphagidae) occur where. Meliphagidae assemblages showed increasing phylogenetic clustering along a gradient of decreasing precipitation away from the ancestral state, with statistically significant clustering observed in the driest sites. Miller et al. (2013) controlled for spatial auto-correlation and found that the significance of the overall relationship was unaffected by such auto-correlation, but the null models they used allowed any species to occur in any quadrat.

To better account for the influence of dispersal limitations, I re-calculated the significance of the observed phylogenetic community structure metrics in Meliphagidae assemblages as compared with expectations from DNM_1_ (both MPD and interspecific MPD). For the matrix describing distances between quadrats, I used: 1) great circle distances calculated with the Haversine formula; 2) the Euclidean distances after a principal components analysis of the 19 bioclim ecological variables (Hijmans et al. 2005), scaled and centered; 3) the product of these two distance matrices. I calculated the significance of the observed metrics against 1,500 randomly assembled matrices with each of the distance matrices. Because results were qualitatively identical with any of the distance matrices, I chose to use the product of the geographic and climate distance matrices and re-ran the analysis with 30,000 matrix randomizations.

## RESULTS

### Statistical behavior and performance of the dispersal null model

With DNM_1_, species’ mean occurrence frequencies across 20 randomized matrices were correlated with their occurrence frequency in the observed CDM (r^2^ = 0.45, p < 0.001). The same was true for their total abundance in the randomized CDMs (r^2^ = 0.79, p < 0.001). With DNM_3_, species’ randomized occurrence frequencies were correlated with observed frequencies (r^2^ = 0.84, p < 0.001), but species’ total randomized abundances were only weakly correlated with observed values (r^2^ = 0.18, p < 0.001).

Averaging across its performance with all tested metrics, DNM_1_ exhibited a 22.8% error rate (15.1% type I, 30.5% type II). However, the bulk of these errors can be attributed to a few metrics (Table 1), namely IAC, *H*_AED_ and, to a lesser extent, PAE and *E*_AED_. With the best performing metric, MPD, the type I and II error rates were 0.1 and 13.8%, respectively. In terms of other metrics that performed reasonably well with DNM_1_, MPD, PD, interspecific MPD, MNTD and AW MNTD all exhibited strong power to detect habitat filtering, but much less power to detect competitive exclusion. The opposite was true of intraspecific MPD and *E*_AED_.

**Table 1.**
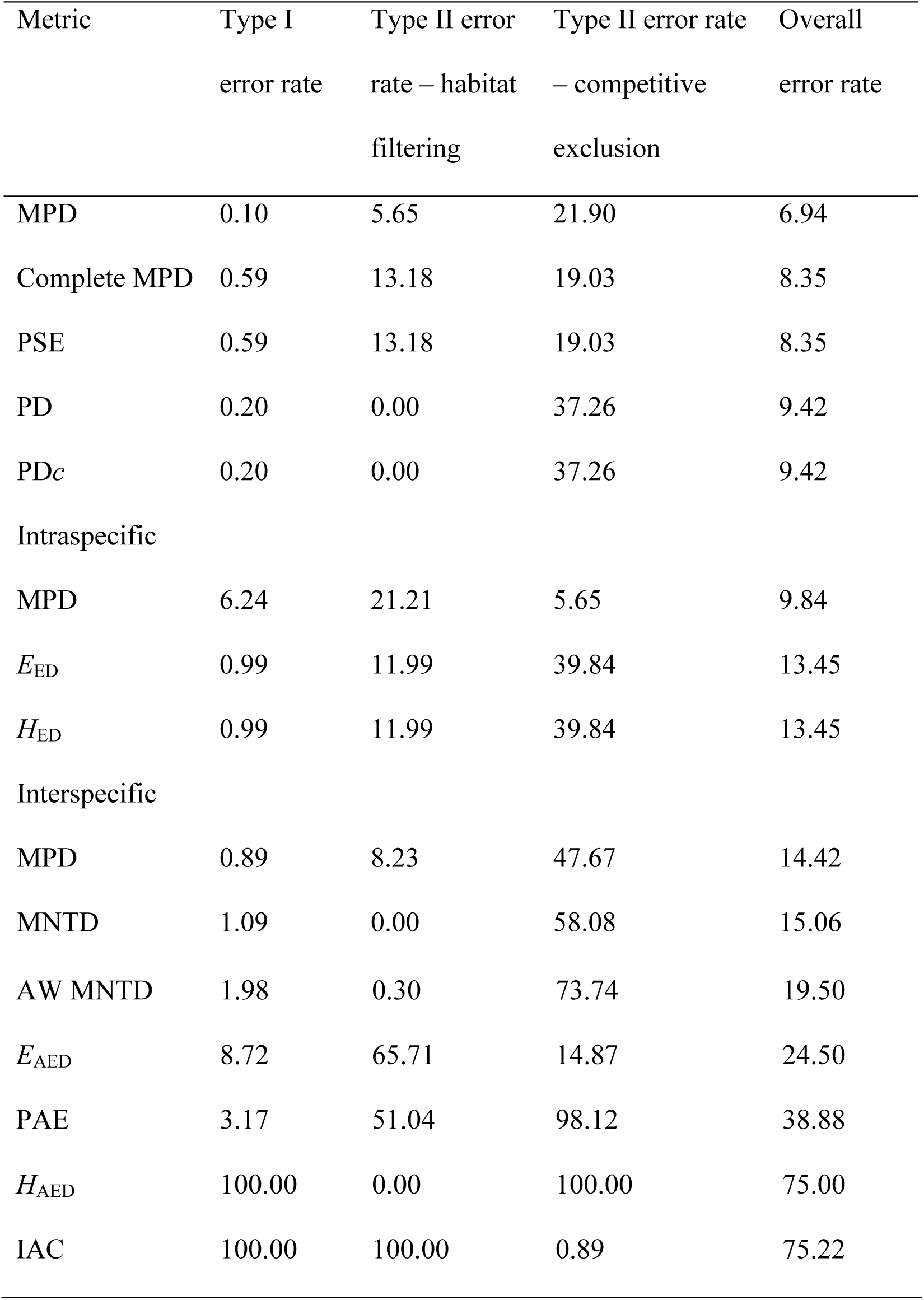
Error rates of the dispersal null model (DNM_1_) with the tested phylogenetic community structure metrics. Type I errors were defined as a randomly assembled community deviating beyond statistical expectations. For example, 0.1% of 1,000 total randomly assembled communities deviated beyond expectations. Type II error rates were defined as communities assembled according to either habitat filtering or competitive exclusion not being considered significantly phylogenetically structured. Metrics are ordered from best- to worst-performing according to the average of the type II and type I error rates.

| Metric | Type I<br>error rate | Type II error<br>rate – habitat<br>filtering | Type II error rate<br>– competitive<br>exclusion | Overall<br>error rate |
| --- | --- | --- | --- | --- |
| MPD | 0.10 | 5.65 | 21.90 | 6.94 |
| Complete MPD | 0.59 | 13.18 | 19.03 | 8.35 |
| PSE | 0.59 | 13.18 | 19.03 | 8.35 |
| PD | 0.20 | 0.00 | 37.26 | 9.42 |
| PD<sub>c</sub> | 0.20 | 0.00 | 37.26 | 9.42 |
| Intraspecific |  |  |  |  |
| MPD | 6.24 | 21.21 | 5.65 | 9.84 |
| $E_{ED}$ | 0.99 | 11.99 | 39.84 | 13.45 |
| $H_{ED}$ | 0.99 | 11.99 | 39.84 | 13.45 |
| Interspecific |  |  |  |  |
| MPD | 0.89 | 8.23 | 47.67 | 14.42 |
| MNTD | 1.09 | 0.00 | 58.08 | 15.06 |
| AW MNTD | 1.98 | 0.30 | 73.74 | 19.50 |
| $E_{\text{AED}}$ | 8.72 | 65.71 | 14.87 | 24.50 |
| PAE | 3.17 | 51.04 | 98.12 | 38.88 |
| $H_{\text{AED}}$ | 100.00 | 0.00 | 100.00 | 75.00 |
| IAC | 100.00 | 100.00 | 0.89 | 75.22 |

DNM_3_ also performed favorably (Table 2). As compared with DNM_1_, it showed increased power to detect simulated community assembly processes, particularly competitive exclusion, with only a slight increase in type I error rates (across all metrics 16.7% type I, 23.2% type II). MPD, again the best performing metric, exhibited an overall error rate of only 2.2%. Both forms of PD also performed well, though they showed less power to detect the effects of competitive exclusion. *H*_ED_ and *E*_ED_, which did not perform well in Miller et al. (2016), performed better than all other tested metrics except MPD and PD.

**Table 2.**
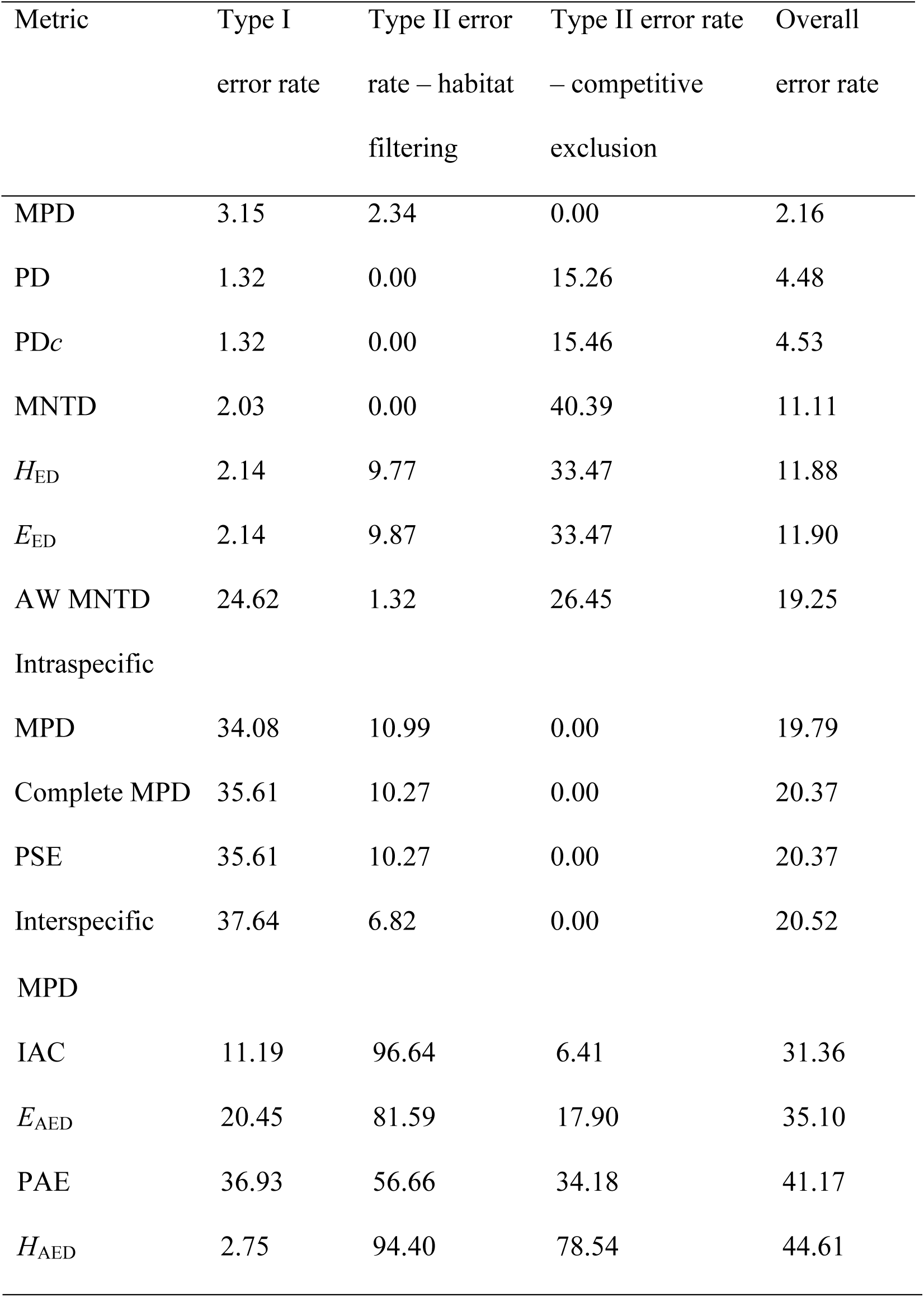
Error rates of the dispersal null model (DNM_3_) with the tested phylogenetic community structure metrics. Type I errors were defined as a randomly assembled community deviating beyond statistical expectations. For example, 0.1% of 1,000 total randomly assembled communities deviated beyond expectations. Type II error rates were defined as communities assembled according to either habitat filtering or competitive exclusion not being considered significantly phylogenetically structured. Metrics are ordered from best- to worst-performing according to the average of the type II and type I error rates.

| Metric | Type I<br>error rate | Type II error<br>rate – habitat<br>filtering | Type II error rate<br>– competitive<br>exclusion | Overall<br>error rate |
| --- | --- | --- | --- | --- |
| MPD | 3.15 | 2.34 | 0.00 | 2.16 |
| PD | 1.32 | 0.00 | 15.26 | 4.48 |
| PD<sub>c</sub> | 1.32 | 0.00 | 15.46 | 4.53 |
| MNTD | 2.03 | 0.00 | 40.39 | 11.11 |
| $H_{ED}$ | 2.14 | 9.77 | 33.47 | 11.88 |
| $E_{ED}$ | 2.14 | 9.87 | 33.47 | 11.90 |
| AW MNTD | 24.62 | 1.32 | 26.45 | 19.25 |
| Intraspecific |  |  |  |  |
| MPD | 34.08 | 10.99 | 0.00 | 19.79 |
| Complete MPD | 35.61 | 10.27 | 0.00 | 20.37 |
| PSE | 35.61 | 10.27 | 0.00 | 20.37 |
| Interspecific | 37.64 | 6.82 | 0.00 | 20.52 |
| MPD |  |  |  |  |
| IAC | 11.19 | 96.64 | 6.41 | 31.36 |
| $E_{\text{AED}}$ | 20.45 | 81.59 | 17.90 | 35.10 |
| PAE | 36.93 | 56.66 | 34.18 | 41.17 |
| $H_{\text{AED}}$ | 2.75 | 94.40 | 78.54 | 44.61 |

### Testing the dispersal null model on an empirical dataset

As in Miller et al. (2013), there was strong signal of increasing phylogenetic clustering along a gradient of decreasing precipitation. For instance, using the product of the climatic and geographic distance matrices, SES MPD and SES interspecific MPD were positively correlated with log_10_ mean annual precipitation (r^2^ = 0.14 and p < 0.001, and r^2^ = 0.41 and p < 0.001, respectively). However, the significance of individual quadrat deviations beyond expectations can and did change with DNM_1_. Only one and two of the quadrats were considered significantly phylogenetically clustered with MPD and interspecific MPD, respectively. Similarly, two and zero quadrats were considered significantly phylogenetically overdispersed with MPD and interspecific MPD, respectively. Results were qualitatively identical with any of the distance matrices (i.e., few individual quadrats deviated beyond statistical expectations). Thus, when compared against a null model that simulates realistic dispersal probabilities, the overall pattern of increasing phylogenetic clustering along a gradient of decreasing precipitation did not change, but few if any individual sites deviated beyond null expectations.

## DISCUSSION

Null models in ecology have been contentious for well over 40 years. Many technical improvements have been developed over this time, including models that account for species-specific patterns of spatial distributions (Roxburgh and Chesson 1998; Roxburgh and Matsuki 1999). A great deal of sound reasoning and guidance has also been offered (Gotelli and Graves 1996; Gotelli 2000; Gotelli and Entsminger 2001; Ulrich and Gotelli 2010). But to my knowledge, no null model that maintains realistic, quadrat-specific dispersal pressures has yet been developed. In this paper I introduced and tested such a dispersal null model (DNM). This is similar to recent efforts to include other biologically important processes in the null model. For instance, Pigot and Etienne (2015) showed that incorporating allopatric speciation into null models erases signatures in phylogenetic community structure that were previously considered to represent competitive exclusion. Indeed, those authors and other recent reviews (Gotelli and Ulrich 2012) highlighted the need for a DNM.

To date, the conceptual link remains weak between neutral models for community assembly and null models for phylogenetic community structure. The DNM and other recent null models provide the foundation for a bridge to link the ideas, but that bridge remains to be built. Future researchers will need to merge ideas of ecological sorting with those of evolutionary processes, e.g. competitive exclusion versus character displacement (or allopatric speciation). Ultimately, a model linking dispersal, speciation and extinction may allow researchers to untangle the influences of these processes in community assembly.

In this paper, the overall error rates of DNM_1_ and DNM_3_ were 22.8% and 19.95%, respectively. In a previous test of null model performance (Miller et al. 2016), across all metrics, the regional model showed the lowest overall error rates (8%), followed by the 3x, 2x, trial swap, independent swap, and frequency concatenated by richness models, which all had overall error rates of approximately 19%. The 1s, richness and frequency concatenated by quadrat models showed error rates of 25-27%. This would suggest the DNM was outperformed by a number of other null models. However, error rates calculated across all metrics are misleading in this case, in that some metrics performed quite well with the DNM, while others performed very poorly. For instance, as compared with the regional null model in Miller et al. (2016), where MPD showed 6.2 and 3.2% type I and II error rates, respectively, with DNM_1_ these rates were 0.1 and 13.8% (Table 1), while with DNM_3_ they were 3.2 and 1.2% (Table 2).

As compared to other null models, the decrease in type I error rates for the DNM is attributable to the fact that simulated quadrats closely resemble observed quadrats in species richness and composition. To deviate beyond expectations, observed quadrats need to show strong signals in terms of co-occurrence and/or, for abundance-weighted metrics, the relative abundances of co-occurring species. As compared with DNM_1_, the increased power of DNM_3_ to detect competitive exclusion seems to be because the latter does not incorporate a species’ relative abundance into its probability of being settled in simulated quadrats. So, if a species is rare in a given quadrat as a function of competition with co-occurring species, this element is randomized in the simulated CDMs, allowing appropriate rejection of the null hypothesis. Conversely, there was a slight overall decrease in power with DNM_3_ to detect habitat filtering. On the surface it would seem this is because if a species’ abundance attenuates away from the center of its distribution, the DNM_3_ may occasionally settle the species at high abundances towards the periphery of its range. In practice, however, most abundance-weighted metrics actually showed increased power to detect habitat filtering with DNM_3_, and the overall decrease in power in this respect as compared with DNM_1_ can be ascribed to *E*_AED_ and *H*_AED_ showing striking decreases in power to detect habitat filtering.

Notably, the non-abundance-weighted metrics *H*_ED_ and *E*_ED_ performed reasonably well with both null models tested here, particularly DNM_3_. These metrics had previously shown poor statistical performance (Miller et al. 2016). Overall, other non-abundance-weighted metrics (MNTD, PD, MPD) also outperformed abundance-weighted forms with DNM_3_. As noted above, abundance-weighted metrics were better than non-abundance-weighted metrics at detecting habitat filtering with DNM_3_, so this overall decrease in abundance-weighted metric performance with DNM_3_ is attributed entirely to a stark increase in their type I error rates (Table 2). Since DNM_3_ maintains species’ occurrence frequencies but not abundances, this increase in type I error rates is to be expected. Abundance-weighted metrics are particularly driven by changes in the abundance of distantly related species. So, for instance, a monotypic genus that occurred regularly but at low abundance in an observed CDM might occur regularly but at high abundance in simulated CDMs, thereby triggering false positives.

The choice of which null model to use cannot be informed by statistical performance alone (Gotelli 2000). The null hypothesis to be tested must inform the choice as well. When used on an empirical dataset where dispersal limitations almost certainly influence probability of arrival at a site (Miller et al. 2013), DNM_1_ did not change previous results that co-occurring Meliphagidae species are more closely related in arid areas. However, the number of quadrats considered significantly phylogenetically clustered was much reduced. Thus, while the overall relationship remains unchanged, the strong signal of Meliphagidae phylogenetic clustering in arid regions would not have been detected had that study focused on single sites and their statistical significance, rather than the slope of unstandardized MPD across climate gradients.

As programmed here, the DNM runs more slowly than previously defined null models, since it requires the use of an indeterminate loop to create each cell in the random matrix, rather than using simple matrix shuffling. Fortunately, the model is available in a multithreaded version, which permits parallel matrix randomizations and a corresponding decrease in total computing time. For instance, constructing 30,000, 696 quadrat by 75 species matrices against which to compare the observed Meliphagidae CDM took ~6 hrs on a MacBook Pro with a 2.5 GHz processor and 16 GB RAM.

The DNM should be applicable to a wide breadth of research questions. There is one situation, however, under which the DNM will fail to run. If any quadrats within an observed community data matrix (CDM) contain more species than are available in the sum of other quadrats, then the model cannot run to completion. For example, if *SR*(*i_O_*) = 15, and the remaining quadrats contain only 12 unique species in total, then the model would loop indefinitely trying to find 15 species to fill *i_R_*. The DNM function has an internal check for this problem, and returns an error message if it is manifest in a CDM. Aside from this hopefully unusual empirical situation, if users are able to generate a distance matrix summarizing dispersal probabilities between quadrats (e.g., geographic or climatic distances between sampling sites), DNM will almost certainly be easy to implement. It strictly maintains species richness, approximately maintains species occurrence frequencies, overall abundances, and realistic dispersal probabilities, and showed suitable performance when compared against simulated community processes.

## ACKNOWLEDGEMENTS

Funding for this study was provided by the National Science Foundation (DBI-1402506). Simulations were run on the Grethor cluster at the University of Missouri, St. Louis, and the Lewis cluster at the University of Missouri, Columbia. Cody Hinchliff, Matthew Pennell, Luke Harmon and three anonymous reviewers provided valuable input during the writing of the manuscript.

